# Unveiling Interaction Signatures Across Viral Pathogens through VASCO: Viral Antigen-Antibody Structural COmplex dataset

**DOI:** 10.1101/2025.03.11.642737

**Authors:** Kenny Wong, Ishaan Subramanian, Emma Stevens, Srirupa Chakraborty

**Affiliations:** Department of Chemical Engineering, Northeastern University, Boston, MA; Department of Bioengineering, Northeastern University, Boston, MA; Department of Chemistry and Chemical Biology, Northeastern University, Boston, MA; Department of Physics, Northeastern University, Boston, MA

## Abstract

Viral antigen-antibody (Ag-Ab) interactions shape immune responses, drive pathogen neutralization, and inform vaccine strategies. Understanding their structural basis is crucial for predicting immune recognition, optimizing immunogen design to induce broadly neutralizing antibodies (bnAbs), and developing antiviral therapeutics. However, curated structural benchmarks for viral Ag-Ab interactions remain scarce. To address this, we present **VASCO** (Viral Antibody-antigen Structural COmplex dataset), a high-resolution, non-redundant collection of ∼1225 viral Ag-Ab complexes sourced from the Protein Data Bank (PDB) and refined via energy minimization. Spanning Coronaviruses, Influenza, Ebola, HIV, and others, VASCO provides a comprehensive structural reference for viral immune recognition. By comparing VASCO against general protein-protein interactions (GPPI), we identify distinct sequence and structural features that define viral Ag-Ab binding. While conventional descriptors show broad similarities across datasets, deeper analyses reveal key sequence-space interactions, secondary structure preferences, and manifold-derived latent features that distinguish viral complexes. These insights highlight the limitations of GPPI-trained predictive models and the need for specialized computational frameworks. VASCO serves as a critical resource for advancing viral immunology, improving predictive modeling, and guiding immunogen design to elicit protective antibody responses. By bridging sequence and structural immunological datasets, VASCO should enable better docking, affinity prediction, and antiviral therapeutic development—key to pandemic preparedness and emerging pathogen response.

## INTRODUCTION

Antigen (Ag) and antibody (Ab) interaction is a fundamental feature of our immune system[1], wherein antibodies evolve to recognize and bind to antigens, forming stable complexes and ultimately perturbing antigenic function and facilitating their neutralization. Beyond their biological significance, antibodies have also become invaluable tools in laboratory settings and are increasingly employed as therapeutic agents[2]. By virtue of their capabilities to recognize antigen epitopes with high specificity[3], and their modular anatomy[4], these antibodies form essential targets for diagnostic and therapeutic mechanisms as well as experimental biology assays[5–7]. The burgeoning field of antibody engineering seeks to understand Ag-Ab binding, stability, and immunogenic properties. Yet this field suffers from expensive resource requirements, biosafety concerns, and research reproducibility crisis[8], making computational methods essential alternatives in this endeavor. While computational tools have proven instrumental in antibody engineering, the reliance on antibody sequence specific data rather than Ag-Ab structural complexes has become a prevailing trend due to greater data availability [9]. To facilitate antibody-related computational analyses, various databases, such as DIGIT[10], IMGT[11], Abysis[12], Antibodypedia[13], and Antibody Registry[14] have been established, each offering distinct advantages in terms of sequence information. However, to get a molecular level understanding of antibody response, it is crucial to consider structural nuances that govern Ag-Ab interactions beyond just the sequence space. Such structural information can provide essential understanding about the conformational topology, dynamics, and binding affinities that underpin this immune response, facilitating rational design of vaccines and therapeutic antibodies.

The growth of publicly available conformational data for Ag-Ab complexes in the Protein Data Bank (PDB)[15] reflects the recognition of the pivotal role of structural information of interactions. The PDB has witnessed a substantial surge in antibody structures, exponentially increasing over the years, and now constitutes approximately 4.2% of the total entries (March 2025). Many contributions have already been made by datasets such as SAbDAb[16], Thera-SAbDAb[17] and IEDB-3D[18]. However, this increasing data availability opens the possibility of exploring underlying patterns in antibody – antigen interaction complexes whichdemands the benchmarked curation of structural data for specific classes of proteinsthat goes beyond sequence or antibody information, but more of a focus on the interfaces of interactions.

A critical subset of Ag-Ab interactions that deserve detailed scrutiny are viral antibody-antigen interaction complexes. The frontline defense in virus-host interactions entails antibodies binding to viral antigens to neutralize their function or recruit other immune components for targeted destruction. In the realm of viral Ag-Ab interactions, resource, safety, and reproducibility challenges are heightened. The dynamics of viral infections[19], coupled with the inherent variability of viral strains[20], necessitate a meticulous understanding of Ag-Ab characteristics where structural characterization of antibody-antigen complexes takes center stage. Antibodies have a typical two-chain modular anatomy[21] where the antigen recognition site is largely limited to the complementarity determining regions (CDRs) that include three hypervariable (HV) loops from each chain. Based on their structural and sequence variability, the antigen binding region (paratope) of antibodies personate their binding specificity[22], with much of the uncertainty coming from these variable loops[23]. Moreover, it has been shown that not all the CDRs may be involved in the interactions in many cases, or some parts of the paratope also fall outside the CDRs, contributing to the intricacy of binding. Identification of viral envelope antigen residues that form the binding interface to antibodies (i.e., the epitope region) is even more challenging due to an apparent lack of common features. While general PPI prediction models have seen success in protein docking and interaction scoring, they consistently struggle when applied to Ab-Ag interactions, yielding lower performance[24, 25].

The need of the hour is to curate and systematically evaluate structural data of viral Ag-Ab interaction complexes along with comparative analyses against general protein-protein interactions (PPI). This approach can shed light on the unique features and challenges posed by viral interactions, offering valuable insights that extend beyond the immediate context of infection. Such a curated dataset will play a pivotal role in advancing viral immunology research by providing a foundation for predictive modeling, therapeutic development, and a deeper understanding of the complexities inherent in viral antibody responses. Understanding structural patterns in viral antibody-antigen interactions is essential for evaluating the accuracy of computational tools and predictive models. Specialized benchmarks, such as DOCK6[26], Docking Benchmark 5.0[27], and Affinity Benchmark versions 1 and 2[27], already provide a standardized platform for assessing the performance of algorithms in general PPIs. In contrast, by establishing benchmarks tailored to the unique features of viral Ag-Ab associations, researchers can refine and improve predictive capabilities specific to attributes of viral responses. Comparative analyses with general PPIs will enable the identification of distinguishing features, such as the sequence-structure topology of CDR loops and the adaptability of antibodies to diverse viral strains. Understanding these distinctions can inform the development of targeted therapies and contribute to our broader understanding of immune responses.

Curated datasets of viral Ag-Ab interaction complexes, evaluated against general protein-protein interactions, will find applications across various scientific research domains. Insights gained from curated datasets can aid in the design of targeted therapies and vaccines against viral infections. By analyzing viral antibody-antigen interactions, researchers can identify conserved structural motifs that may be crucial for vaccine efficacy. Understanding the structural patterns of effective antibody responses can also inform the development of antiviral drugs and prophylactic measures[28, 29]. The insights derived from these comparative analyses can contribute to the identification of potential biomarkers and therapeutic targets. Structural patterns derived from curated datasets can further contribute to the modeling and prediction of viral diseases and their evolution.

In this paper, we present VASCO (Viral Antibody-antigen Structural COmplex dataset), a curated collection of 1225 high-resolution, non-redundant viral antigen-antibody (Ag-Ab) interaction complexes in an energy-relaxed conformation close to their crystal structure local minimum, with resolutions better than 5 Å. The dataset encompasses a diverse range of viral species, including SARS-CoV-2, Influenza, Ebola, and HIV, providing a comprehensive structural reference for viral immune recognition. Additionally, VASCO includes a detailed set of structural and physicochemical features relevant for training, validation, and testing of machine learning models, facilitating predictive modeling of antibody-antigen interactions. To assess the distinct properties of viral Ag-Ab interactions, we compared VASCO against two control datasets of general protein-protein interactions (GPPI), comprising 2000 heterodimeric and homodimeric complexes. While conventional structural features—such as contact surface area, hydrogen bonding, and non-covalent interactions—showed broad similarities between datasets, deeper analyses uncovered key sequence-structure signatures and manifold-derived latent features that distinguish viral Ag-Ab interfaces. These findings underscore the limitations of existing GPPI-based predictive models in capturing the complexities of Ag-Ab binding and emphasize the need for specialized computational frameworks tailored to antibody-antigen interactions. By providing a benchmark dataset and structural insights into viral immune recognition, VASCO serves as a valuable resource for advancing viral antibody engineering, vaccine design, and computational immunology.

## RESULTS

### Constructing VASCO as a Dataset to Bridge the Gap in Ab-Ag Interaction Modeling

The VASCO dataset was curated through an exhaustive search of the Protein Data Bank (PDB) by querying its API for structural data of viral antigen-antibody (Ag-Ab) complexes. We focused on high-resolution complexes, selecting those with a resolution better than 5Å, which ensures structural reliability for downstream analyses. The dataset comprises 1225 non-redundant viral antibody-antigen interaction complexes. To ensure structural consistency and reduce artifacts from crystallographic, cryo-EM, or NMR-derived conformations, all structures in VASCO underwent local energy minimization, preserving their native-like binding states while resolving steric clashes or structural irregularities. The viral species represented in the dataset were further categorized as follows (See **Figure 1**): SARS-CoV and CoV-2, MERS and related coronaviruses: 544 structures; HIV: 183 structures; Influenza : 144 structures; Ebola: 26 structures; other miscellaneous viral Ag-Ab complexes: 103 structures. This dataset, referred to as VASCO (Viral Antibody antigen Structural COmplex dataset), aims to support predictive modeling efforts for antibody-antigen interactions, which are distinct from general protein-protein interactions (GPPI) due to the structural complexity of antibody binding interfaces. In particular, antibody-antigen binding sites exhibit highly variable topologies due to the complementarity-determining regions (CDRs) and variable loops, making accurate predictions challenging using models developed for more general protein-protein interactions. To highlight these differences, we complemented the VASCO dataset with two control datasets of GPPI interfaces: heterodimers and homodimers, each comprising 2000 randomly selected interactions from the PDB, similarly filtered for structural resolution below 5 Å. These general protein-protein interaction datasets (GPPI) provide a basis for comparison to help assess the unique structural properties of viral antibody-antigen interactions. The VASCO dataset includes a range of typical protein interface descriptors such as contact surface areas, hydrogen bond counts, and secondary structure involvement, which are standard in PPI classification. However, initial comparisons between VASCO and the GPPI datasets showed no significant deviations in most structural features across the datasets. Then the obvious question becomes, how do the current state of the art GPPI interaction prediction methods fail to perform as well Ab-Ag interaction prediction? Interestingly, deeper analysis revealed notable differences in specific feature distributions: (i) Sequence space of the contact residues: Antibody-antigen interfaces showed distinct patterns in sequence composition at the contact points compared to GPPI. (ii) Secondary structure features: Certain differences emerged in the involvement of secondary structural elements at the interfaces, with viral antibody-antigen interactions displaying more variability. (iii) Manifold-derived latent features: Using dimensionality reduction techniques, we identified latent features—complex representations derived from structural manifolds—that exhibited significant differences between VASCO and GPPI datasets. These findings indicate that while many general protein interface features overlap between viral Ag-Ab interactions and GPPI, specific characteristics, especially in the sequence and structural context, set antibody-antigen interactions apart, underscoring the need for specialized predictive models for these complexes.

**Figure 1:**
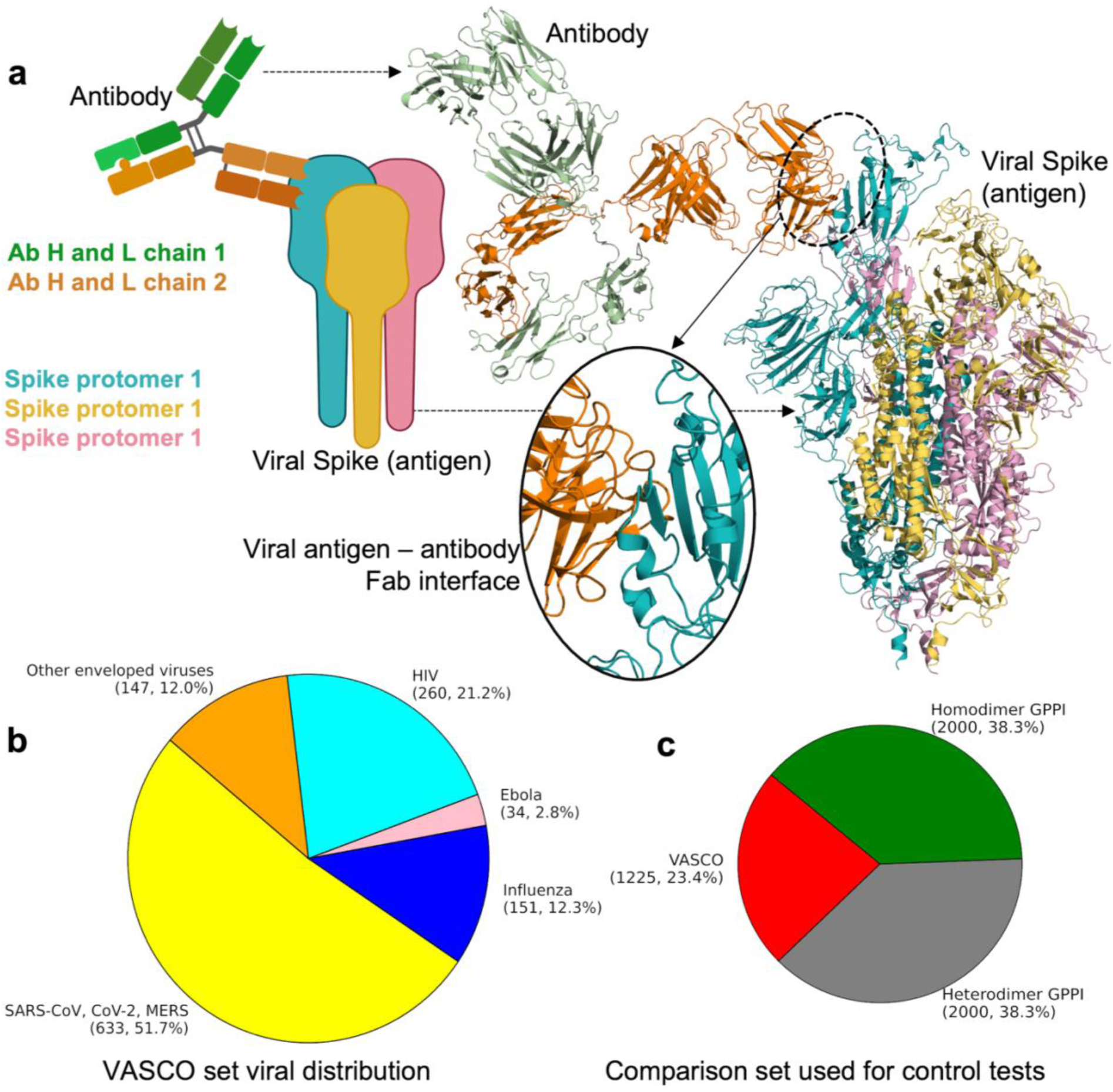
The VASCO set of energy-minimized viral antigen-antibody structural complexes. (a) Schematic and actual structural representations of viral antigens interacting with antibody Fab domains, with a zoom-in on the example contact interface of SARS-CoV2 spike interacting with H014 Fab (PDB 7CAC). The immunoglobulin full structure was aligned by the fab domain for representation purposes (PDB 1IGY). (b) Composition of the VASCO dataset. (c) Composition of the comparison set of viral Ag-Ab with general PPI complexes.

### Distinctive Amino Acid Contact Profiles Define Viral Antibody-Antigen Interface Signatures

**Figure 2** presents the sequence contact maps comparing the frequency of amino acid contacts between viral antibody-antigen (Ab-Ag) interfaces and general protein-protein interactions (GPPI) from homodimer and heterodimer control sets. A striking observation is that while the contact maps for the homo and hetero control sets are nearly indistinguishable (**Figure 2b,c**), the viral Ab-Ag contact profiles exhibit clear and distinct patterns. In the viral Ab-Ag interfaces, serine (Ser) and tyrosine (Tyr) residues from the antibodies consistently form high-frequency contacts with a variety of antigen residues. Additionally, glycine (Gly) from the antibody shows a high occurrence of contacts with asparagine (Asn), tyrosine (Tyr), and glycine (Gly) on the antigen surface (**Figure 2a**). The significance of Tyr, Ser and Gly on Ab CDRs have been widely noted in literature [30–32]. Among charged residues, the Lys-Asp (lysine from antigen and aspartic acid from antibody) pair is prominent, standing out as a frequent contact. In contrast, the control sets show a much more diverse range of contact pairs, with arginine (Arg) and leucine (Leu) dominating the profile by making frequent contacts with several amino acids, followed by Arg-Phe (phenylalanine) pairs from both sides of the protein interface. Interestingly, in the HIV Ab-Ag interfaces, the serine population diminishes while glycine contacts from both the antibody and antigen sides increase significantly (**Figure 2d**). Glycine mutations have been known to increase the potency and breadth of HIV-1 broadly neutralizing antibodies[33, 34].Despite this shift, the overall contact profiles remain fairly consistent across the different viral species, underscoring the unique signature of viral antibody-antigen interactions. This distinct contact profile, dominated by Ser, Tyr, and Gly from antibodies interacting with specific antigen residues, highlights the unique structural features of viral Ab-Ag interfaces. The recurrence of these specific residue interactions across viral species suggests that pairwise contact frequency signatures encode critical information about antibody recognition. These patterns should be explicitly incorporated into predictive modeling approaches, as they may offer a more biologically relevant and interpretable basis for scoring and ranking antibody-antigen interactions.

**Figure 2:**
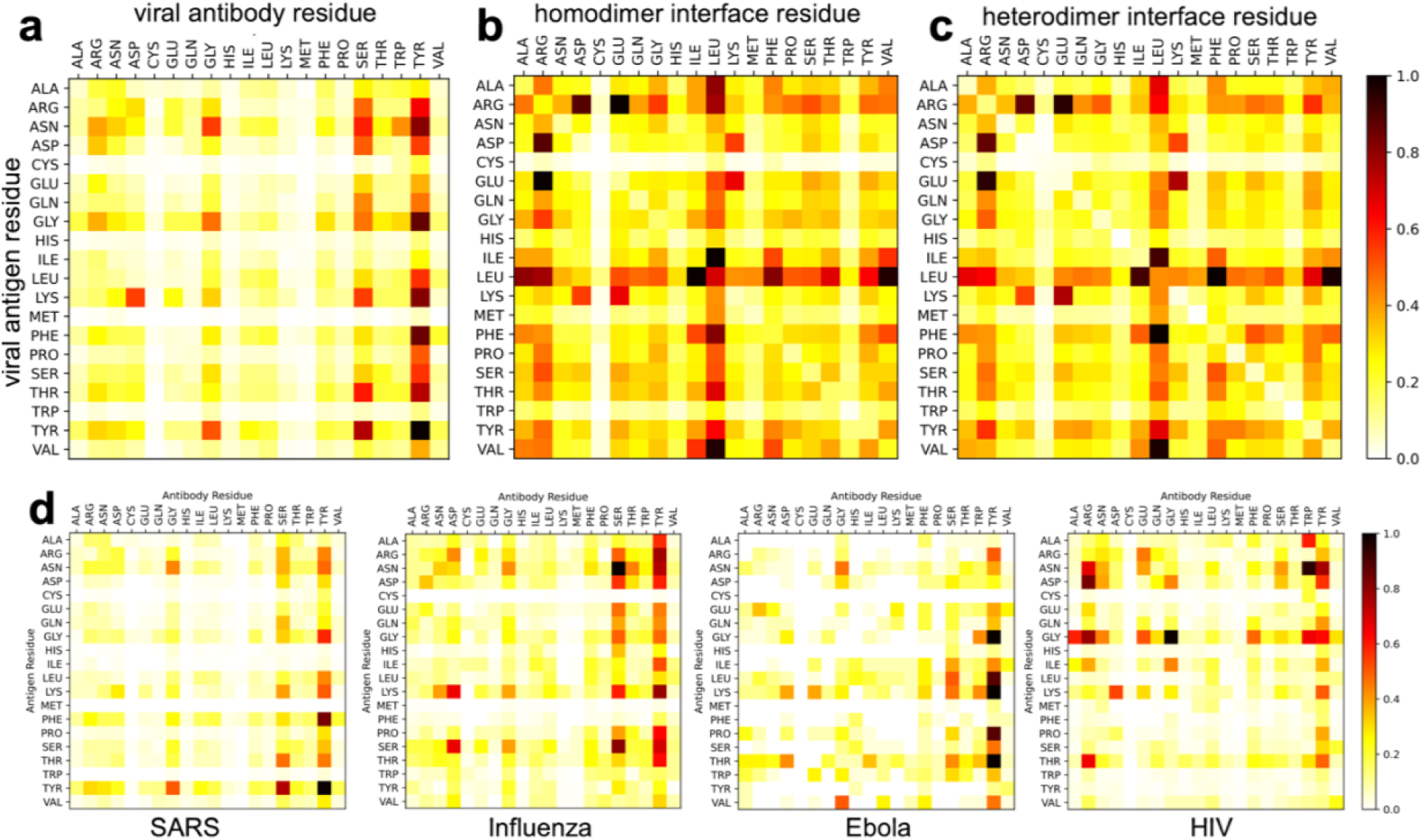
Pairwise contact profile between different amino acids at interaction interfaces. (a) Sequence contact pair frequencies at viral Ag-Ab interface, (b) homodimer, and (c) heterodimer interaction surfaces. (d) Virus-specific sequence contact pair profile.

### Ab-Ag Interactions Share Interface Properties with General PPIs but Highlight the Need for Specialized Models

To investigate the structural characteristics of the viral antibody-antigen (Ab-Ag) interactions in the VASCO dataset, we compared the contact surface area (CSA), number of inter-residue non-covalent contacts, hydrogen bonds, and intermolecular contacts per unit area with those of general protein-protein interactions (GPPI) from homodimer and heterodimer datasets. These comparisons are illustrated in **Figure 3**, with panels showing the results for each feature. In **Figure 3a**, we compare the CSA across the full VASCO set, individual viral species (SARS, Influenza, Ebola, HIV), and the GPPI control sets. While the median CSA is not significantly different between the viral Ab-Ag complexes and the GPPI datasets, the variability is noticeably greater in the control sets. This broader spread of CSA values in the homodimer and heterodimer datasets reflects more diverse surface area exposure in general PPIs, whereas viral antibody-antigen interactions are more uniform in terms of accessible surface area. This is not very surprising, given the strong structural similarities between different Fabs. **Figure 3b** presents the number of non-covalent contacts between the binding molecules. While the median number of contacts is consistent across all groups, the quartile ranges is again significantly larger for the homodimer and heterodimer datasets, indicating that general protein-protein interfaces exhibit greater variability in the number of non-covalent contacts compared to the more constrained viral Ab-Ag interactions. **Figure 3c** presents the number of hydrogen bonds at the interfaces. Here too, the median values are comparable across the viral and GPPI datasets, with significantly larger lower-to-upper quartile spread in the GPPI. Lastly, in **Figure 3d**, we compare the average presence of contacts per unit area of interaction. Unlike the other three measures, here the variabilities are comparable between the VASCO and the control GPPI sets. Similarly, H-bonds per unit area of contact also remained similar between different viral Ag-Ab and general proteins (**Supplementary Figure S1a**). These findings indicate that commonly used interface descriptors—contact surface area, non-covalent interactions, and hydrogen bonds—are not fundamentally different between viral Ag-Ab interactions and general PPIs. Instead, they largely scale with the available interaction surface area. Given this similarity, the consistently poorer performance of ML-based docking and scoring models on Ab-Ag complexes remains an intriguing challenge. This suggests that the key distinguishing factors for Ab-Ag interactions may lie beyond these traditional structural features, underscoring the need for specialized predictive models and alternative feature selection strategies tailored to the unique constraints of antibody-antigen recognition.

**Figure 3:**
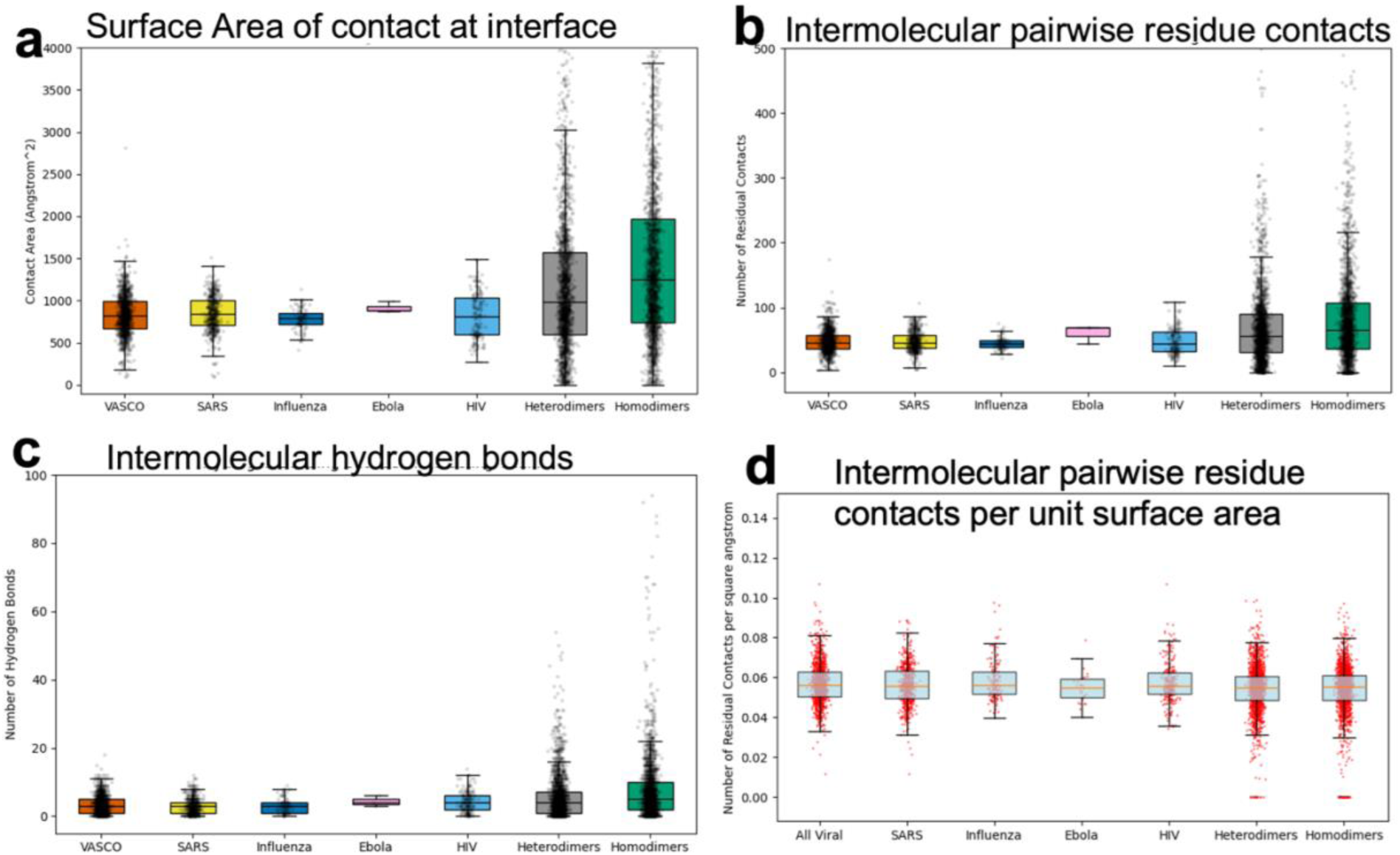
Structural contact features compared between VASCO and control general inter-protein interactions. (a) Median and quartile ranges of surface area (angstrom-squared) of contact between proteins. (b) Median and quartile ranges of intermolecular residue-residue non-covalent contacts. (c) Intermolecular hydrogen bond count. (d) Fractional count of inter-residue contacts between participating proteins, per unit surface area of contact.

### β-Strands and Turns Dominate Viral Ag-Ab Interfaces Over Helical Content

**Figure 4** illustrates the secondary structure composition of the viral antibody-antigen (Ab-Ag) interfaces in the VASCO dataset compared to the general protein-protein interaction (GPPI) control sets. Our analysis reveals a higher propensity for β-bridges and turns within the viral interfaces (**Figure 4a**), while showing significantly lower helical content in comparison to both homodimer and heterodimer datasets (**Figure 4b,c)**. Both homodimeric and heterodimeric protein interactions on the other hand have helices as the main structural features at interfaces, also reported earlier[35]. The comparative data indicate that the viral Ab-Ag interfaces favor β-strands and turns, which may contribute to the stability and specificity of these interactions. This is in stark contrast to the GPPI control sets, where a more balanced distribution of secondary structures is observed, characterized by a greater prevalence of helical content. The higher beta-bridge occurrence stems from the viral antigen surface, not from the antibody, which mainly contacts through the disordered CDRs. Notably, the control sets exhibit similar values across different secondary structure distributions, reinforcing the uniformity of their structural characteristics, independent of homomeric or heteromeric composition. Interestingly, the Ebola interactions display a unique profile, with helical content values that fall within the range observed in the GPPI control sets (**Supplementary Figure S1b**). This suggests that the Ebola virus may possess distinct structural features at its Ab-Ag interfaces, potentially influencing its immune evasion strategies and the design of therapeutic antibodies. Overall, these findings highlight the specific structural adaptations present in viral antibody-antigen interactions, marked by an increased occurrence of β-strands and turns alongside reduced helical content compared to general protein-protein interactions. This distinct secondary structure composition is critical for understanding the dynamics and mechanisms underlying viral immune responses.

**Figure 4:**
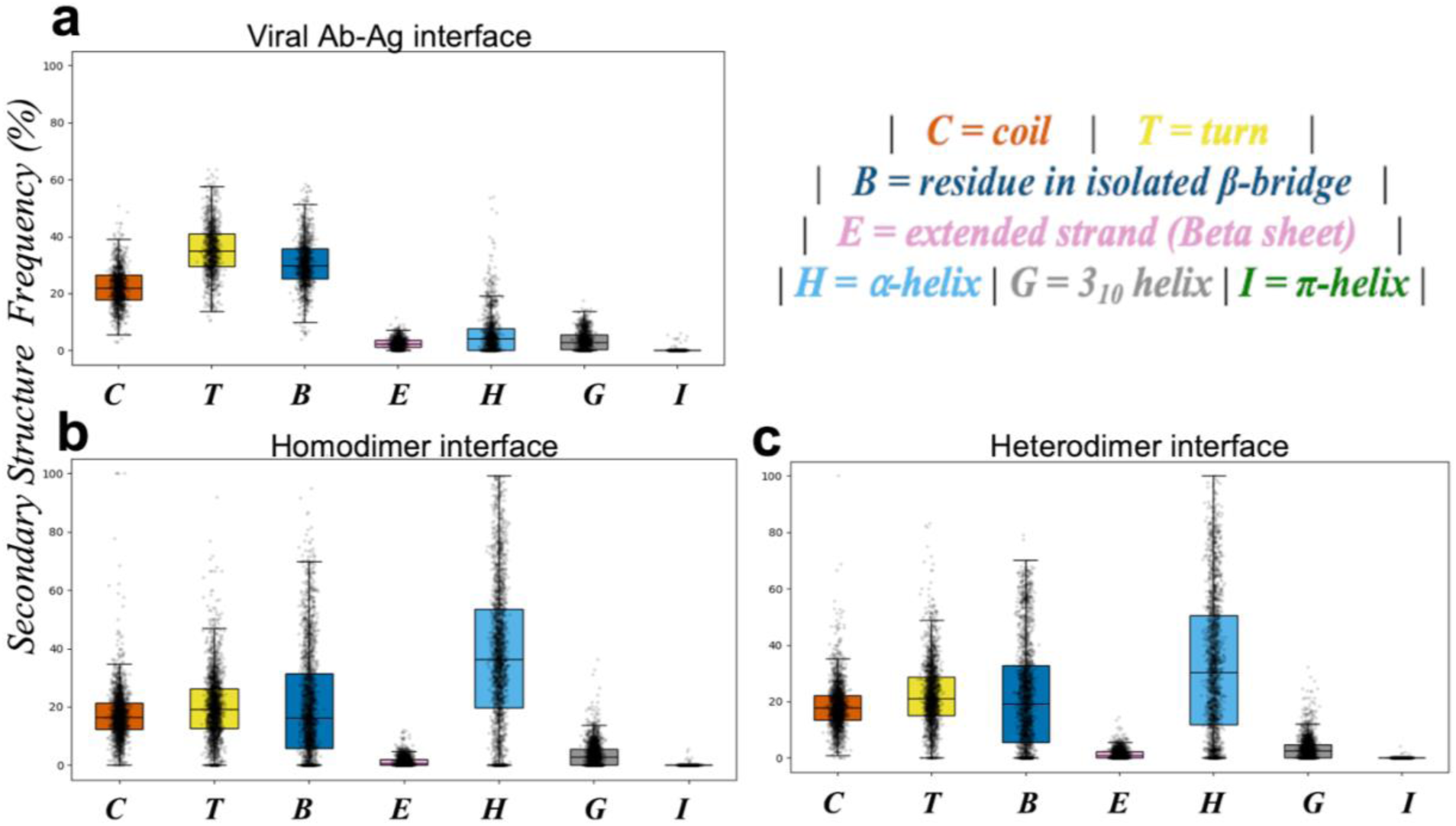
Secondary structure distribution at protein interaction interfaces. (a) Median and quartile ranges of secondary structure distributions over interface residues within the VASCO set. (b) Median and quartile ranges of secondary structure distributions over interface residues within control homodimer and (c) heterodimer interaction interfaces.

### Theoretical Interaction Energies Reveal Stability Trends between Viral and General Interfaces

Interaction energies and affinities can be highly informative for understanding the strength and specificity of antibody-antigen interactions, providing quantitative measures that can guide predictive modeling, improve docking accuracy, and aid in the design of high-affinity therapeutic antibodies. Although 1225 Ab-Ag structures were curated, acquiring corresponding experimental binding affinities or energies proved challenging. We could find less than 5% of these complexes that had experimentally determined binding energies available in the literature, demonstrating the difficulty of obtaining standardized, comparable energy values for such interactions. Given the objective of this study—to provide a comprehensive dataset for data-driven Ab-Ag interaction prediction—it is crucial to include a theoretical measure of comparative binding stability. Despite its limitations in predicting absolute energy values, Molecular Mechanics Poisson-Boltzmann Surface Area (MMPBSA) method is effective in estimating relative binding energy differences, making it suitable for this analysis. **Figure 5** presents the results of theoretically calculated binding free energies for antibody-antigen (Ab-Ag) complexes in the VASCO dataset, using implicit solvent MMPBSA energy calculations. The calculated binding energies were subsequently scaled based on the available experimental data (∼5% of the dataset) to ensure a reasonable comparison. As observed in several other structural features in this study, the trends in binding energies between the VASCO dataset and the control sets (homodimer and heterodimer protein-protein interactions) are within standard deviations of each other. However, there are several noteworthy observations. The median binding energies of viral Ag-Ab complexes fall between those of homomeric and heteromeric protein-protein interactions, with homodimers being the least tightly bound (**Figure 5**, top left). Electrostatic interactions contribute the most to binding stabilization in both VASCO and heteromeric complexes. In contrast, as expected, van der Waals forces serve as the primary stabilizing factor in homomeric interactions but play a comparatively smaller role in viral Ag-Ab binding stability. Additionally, viral complexes exhibit distinct solvation energy characteristics (**Figure 5**, bottom row) —polar solvation energies are less destabilizing, while nonpolar solvation energies provide less stabilization compared to general PPIs.

**Figure 5:**
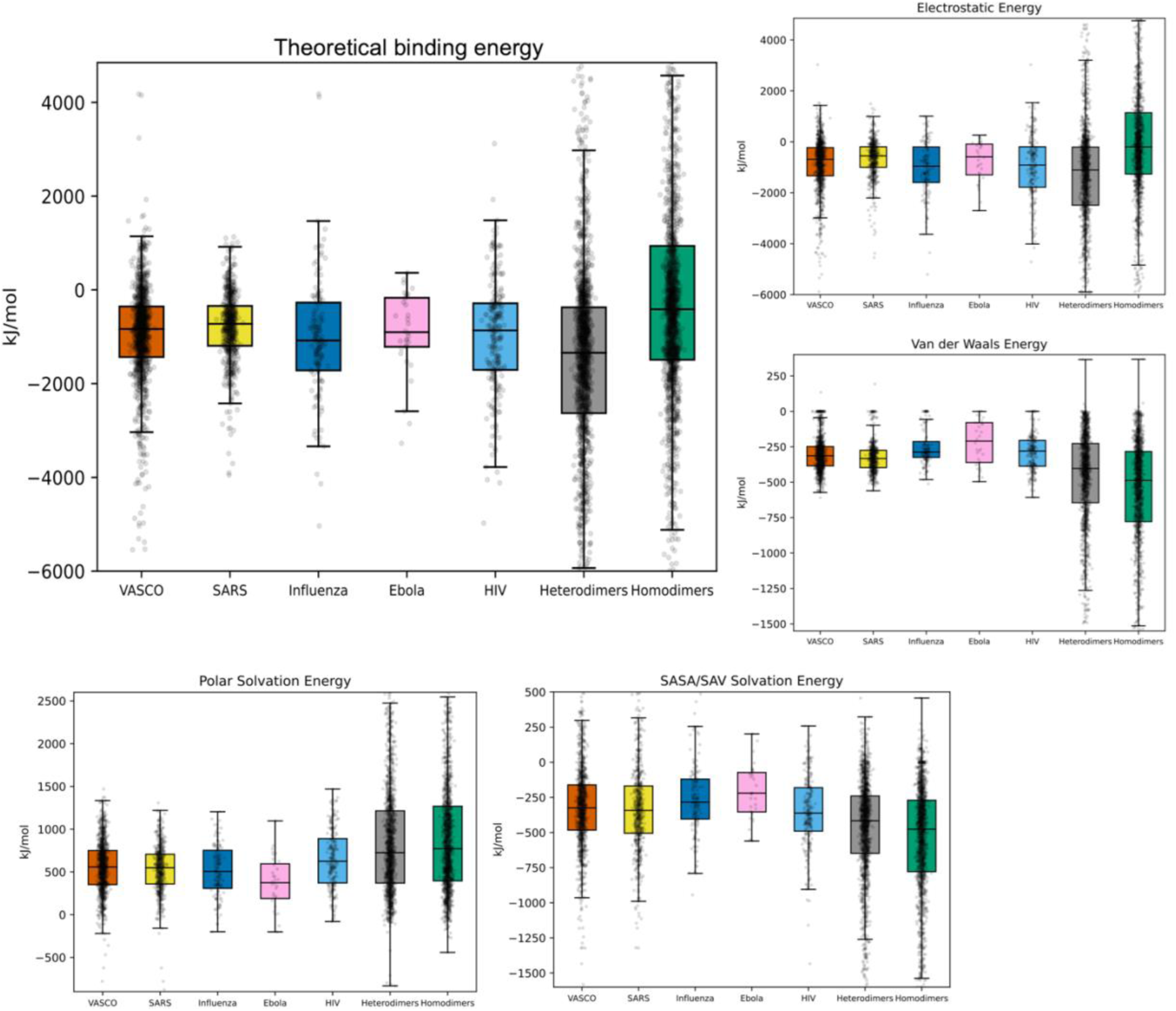
Theoretical binding energy comparison between viral Ag-Ab and general PPIs. The median and quartile ranges are shown. Total binding interaction energy, along with decomposition into electrostatics, van der Waals, and solvation (polar/non-polar) energies are represented in different panels.

One notable observation is the larger variation in binding energies within the control sets, which exhibit a broader range of contact surfaces and interactions. In some cases, particularly in the control sets, energies even show positive values, indicating unstable interactions. These instances highlight the necessity of caution when using experimentally resolved structures (from X-ray crystallography, NMR, or cryoEM) for predictive modeling, as some structures may represent suboptimal, non-stabilized conformations. All energy calculations were conducted following extensive local structure minimizations using the CHARMM36m force field, ensuring accurate theoretical predictions. However, this analysis emphasizes the need for careful treatment of structural and energetic data in any predictive or scoring model, as some structures deviate from their most stable orientations.

### Manifold Learning Reveals Hidden Differences Between VASCO and GPPI interfaces that otherwise remain elusive

Despite limited distinctions found using conventional structural features to describe antibody-antigen (Ab-Ag) interactions, these complexes continue to underperform in general protein-protein interaction (GPPI) binding predictions [REF]. This discrepancy suggests that critical variations in interface architecture may remain hidden in higher-dimensional space, necessitating a more nuanced analysis. To investigate this, we applied several dimensionality reduction techniques to explore patterns in the data that traditional handcrafted descriptors may miss (**Figure 6** and **Supplementary Figure S2**). Principal component analysis (PCA), one of the most commonly used dimension reduction techniques[36], failed to capture any significant differences between the viral Ab-Ag interfaces and the GPPI control sets (**Supplementary Figure S2a**). This lack of distinction was further echoed when we employed nonlinear dimensionality reduction technique of t-distributed stochastic neighbor embedding (t-SNE)[37] (**Supplementary Figure S2b**). Neither method could meaningfully separate viral interactions from the homo and hetero GPPI controls in the reduced feature space. However, employing the isomap method[38] of checking geodesic distances rather than flat Euclidian variations between datapoints, we observed clear distinctions in the architecture of Ab-Ag interfaces. This was further verified by spectral embedding of the manifold[39]. In both methods, the top two embedding coordinates revealed that viral Ab-Ag interactions form a distinctive cluster, markedly different from those of GPPI controls (**Figure 6a, 6b**). This suggests that these nonlinear techniques can capture subtle variations in the interface that remain elusive in linear and conventional approaches. Interestingly, within the VASCO dataset, HIV Ab-Ag complexes also formed a distinct cluster (**Figure 6c,d**). This observation aligns with findings from the sequence contact map analysis (**Figure 4**), further reinforcing the uniqueness of HIV interactions relative to other viral complexes. Based on these manifold-driven latent features, we developed a probability map in the spectral embedding space, spanned by the top two coordinates. This map assigns a negative log-probability score to any interface, providing a measure as to whether the interface likely belongs to an Ab-Ag interaction (**Figure 6e**) or a GPPI control (**Supplementary Figure S2c**), or even the specific subset of HIV interfaces (**Figure 6f**). This classification tool can be applied to novel protein interfaces, offering a powerful way to categorize them based on structural likelihood. Thus, by employing nonlinear dimensionality reduction techniques, we uncovered critical differences in viral Ab-Ag interface architecture that traditional hand-picked descriptors and linear methods failed to detect. These findings underscore the value of manifold learning as a feature selection tool, capable of capturing complex structural patterns that are crucial for Ab-Ag interaction prediction.

**Figure 6:**
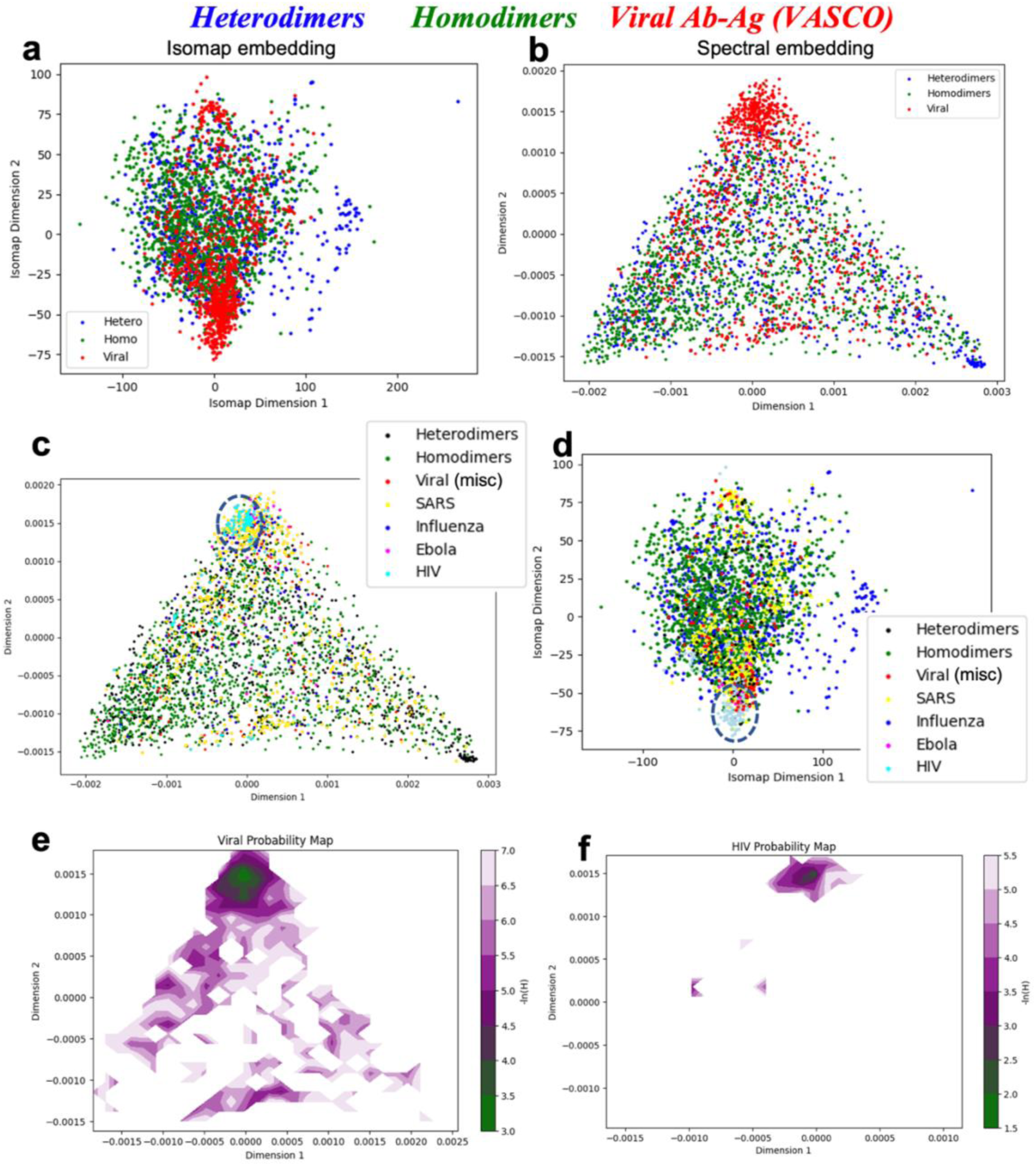
Manifold reduction helps to quantify intrinsic differences between viral Ag-Ab and general PPIs. (a) Isomap and (b) Spectral Embedding of the combined set of VASCO, and general protein-protein interface structures. (c) Spectral embedding and (d) isomap projections of different viral types reveal that HIV Ag-Ab interfaces (cyan) cluster together to form a distinct subclass – shown by circles. (e) Negative log probability map of an interface, measuring its likelihood of being a viral Ag-Ab interface or, specifically, (f) a HIV interface.

### Spectral Embedding Captures Distinct Structural Modes of Viral Ab-Ag Interfaces

To further explore the distinctions uncovered by spectral embedding, we extracted the most representative structures from the three extreme corners in the triangular manifold projection of the spectral embedding space. This allowed us to visualize how the Ab-Ag interfaces from the VASCO dataset vary along the most informative dimensions (**Figure 7**). These representative structures were selected based on their proximity to the mean conformation (by RMSD) within each extreme region of the embedding space (**Figure 7a**), providing insights into the structural diversity across viral interfaces. Among these three extremes, the top cluster **(Set-T)** is the most populated, containing 50% of the VASCO dataset, while the bottom left cluster **(Set-L)** comprises 15% of the interfaces. The bottom right cluster **(Set-R)** is sparsely populated, representing only 3% of the dataset, with the remaining structures distributed throughout the embedded space. Examining the structural differences between these clusters, we find that the Fab architecture remains largely consistent, with the primary variations occurring at the antibody-antigen interface (**Figure 7b**). The CDR loops in Set-T and Set-L exhibit similar alignments, though Set-T shows a slightly more splayed-out heavy chain conformation. In contrast, the Set-R interfaces display a more disordered loop arrangement, with CDR loops protruding outward more prominently.

**Figure 7.**
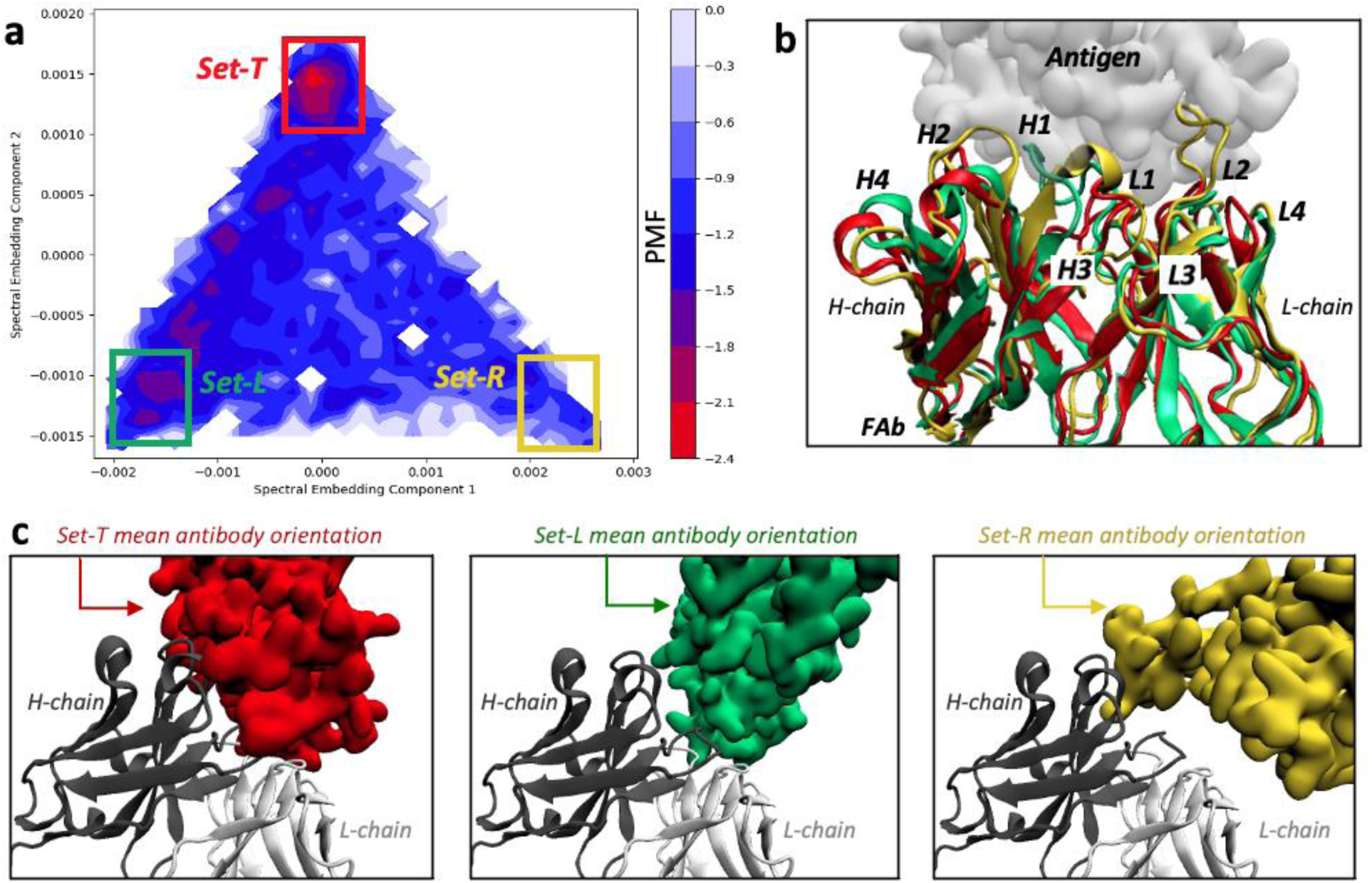
Structural variations in viral Ab-Ag interfaces across spectral embedding extremes. (a) Triangular spectral embedding projection of the VASCO dataset, colored by a weighted histogram PMF representation. More red regions indicate higher population sites. The three extreme clusters (Set-T, Set-L, and Set-R) are marked and colored red, green and yellow respectively. Representative structures were selected based on proximity to the cluster mean by RMSD. (b) Superposition of representative antibody structures from each cluster, illustrating differences in CDR loop conformations and interface orientations. (c) Antigen binding modes across the three clusters, showing differences in surface of contact and chain preference (balanced in Set-T, L-chain biased in Set-L, and H-chain dominant in Set-R).

The nature of antigen contact varies significantly across these clusters (**Figure 7c**). In Set-T, the antigen surface engages both the heavy (H) and light (L) chain CDR loops equally, forming a large surface area of interaction. In Set-L, the contact area is relatively smaller and is slightly biased toward the light chain of the Fab. Meanwhile, in Set-R, which is the least populated conformational state, the antigen makes minimal contact, predominantly with the heavy chain, leading to a significantly reduced binding interface. These findings suggest that while the majority of viral Ab-Ag interactions favor stable, broad-contact conformations (as seen in Set-T), a subset adopts more asymmetric or sparse interactions (Set-L and Set-R). The relatively low occurrence of Set-R structures highlights the instability or rarity of such binding modes, reinforcing the importance of interface-specific constraints in viral antibody recognition. This analysis demonstrates that spectral embedding effectively captures structural heterogeneity in antibody-antigen interfaces, offering a useful framework for categorizing binding modes that could inform predictive modeling and antibody design.

## DISCUSSION

In this study, we have presented VASCO, a curated dataset of viral antigen-antibody (Ag-Ab) structural complexes, comprising over 1225 non-redundant structures that have been energy-minimized to resolve steric clashes and improve geometric consistency. This dataset spans multiple viral families, including SARS, Influenza, Ebola, and HIV, and captures high-resolution structural details of antibody interactions with viral antigens. To contextualize these interactions, we compare the dataset against general protein-protein interaction (GPPI) control sets, including heterodimeric and homodimeric protein complexes. VASCO provides a comprehensive structural representation of human antibody interaction with viruses, offering insights into antibody recognition patterns, viral epitope characteristics, and motifs unique to viral Ag-Ab binding. By integrating traditional structural descriptors with advanced manifold techniques, this dataset serves as a valuable resource for studying antibody-antigen interactions and developing predictive models for viral antibody engineering and therapeutic design.

Our study reveals that while many conventional descriptors do not show significant differences between Ab-Ag and GPPI interfaces, key latent features extracted through nonlinear dimensionality reduction (e.g., isomap and spectral embedding) clearly differentiate the viral complexes. These findings suggest that these manifold embeddings should be used to complement the traditional descriptors in designing a predictive model for Ab-Ag interfaces to attain superior performance from existing GPPI scoring methods.

The curated dataset presented here holds significant implications for advancing the field of antibody engineering and pandemic preparedness. While the predominant contribution to VASCO comes from SARS-related antibodies, this does not limit the dataset’s applicability, as key sequence-structure features remain largely consistent across different viral families, as demonstrated in our comparative analyses. The shared binding properties of viral Ag-Ab interactions, including contact residue compositions and secondary structure preferences, suggest that SARS-CoV-2 serves as a representative model for studying antibody-antigen interactions more broadly. Moreover, the recent COVID-19 pandemic has significantly expanded the availability of high-resolution SARS-CoV-2 antibody-antigen complexes, enriching the dataset and allowing for deeper structural insights that extend to other viral systems.

By providing detailed insights into the structural determinants of viral antibody-antigen interactions, this dataset can serve as a vital resource for researchers working on therapeutic antibody design. Structural differences captured in this dataset will help in improving computational models for predicting antibody binding, guiding rational antibody design with enhanced specificity and binding affinity. The VASCO dataset specifically focuses on viral Ab-Ag interactions, a critical component for understanding immune responses during viral infections. In the context of pandemics, such as COVID-19, studying how antibodies bind to viral antigens can enhance our ability to predict the course of infections and to design therapeutic interventions, including vaccines and monoclonal antibody therapies. This dataset provides the structural foundation for evaluating engineered antibodies and optimizing their therapeutic potential. Given the global relevance of antibody design in combating viral threats, this dataset can inform the development of more effective engineered antibodies. By analyzing the structural features of successful antibody interactions with viral epitopes, this resource will enable the engineering of antibodies with improved binding affinities and therapeutic potential.

While computational techniques such as docking algorithms and binding affinity predictions are essential tools for studying protein-protein interactions, the specific challenges of modeling viral Ab-Ag interfaces require tailored approaches. Benchmarks like Docking Benchmark 5.0, Affinity Benchmark 1 and 2, and DOCKGROUND have provided useful platforms for testing computational models, but the representation of viral Ab-Ag complexes in these benchmarks remains limited. Our dataset aims to bridge this gap by providing a viral-specific benchmark for evaluating the performance of docking and affinity prediction algorithms. Some datasets have been developed for studying antibody-antigen (Ab-Ag) interactions, though they are not specifically focused on viral Ab-Ag complexes. Notable examples include the PECAN dataset[40], which consists of 460 structures, partitioned into 205 for training, 103 for validation, and 152 for testing; the PARAGRAPH dataset[41], containing 1,086 complexes, with 60% allocated for training, 20% for validation, and 20% for testing; and the MIPE dataset[42], comprising 626 structures, with 90% used for five-fold cross-validation and 10% reserved for testing. While these datasets provide valuable benchmarks for Ab-Ag modeling, they encompass a broad range of antigen types, including non-viral targets. In contrast, VASCO is specifically tailored to viral Ab-Ag interactions, capturing the unique structural characteristics of antibodies binding to viral antigens. This specialization makes it a crucial resource for improving predictive models in the context of viral immunity and therapeutic antibody design. The diversity and structural uniqueness of viral Ab-Ag interfaces pose significant challenges for current computational methods. By incorporating the variability and dynamics of viral Ab-Ag complexes into predictive models, this dataset has the potential to improve the accuracy of antibody design and virus neutralization predictions.

One of the key obstacles in Ab-Ag interaction modeling is the structural flexibility of CDR loops, which play a central role in antigen recognition. Viral antibodies often exhibit a higher degree of structural flexibility to adapt to the diverse and dynamic nature of viral antigens. Understanding and incorporating these dynamics into predictive models is critical for capturing the full range of possible interactions. As a result, docking and scoring methods trained on general PPI datasets fail to capture the key determinants of antibody binding, leading to substantial performance gaps. A review of docking studies, including RosettaDock[43], ZDOCK[44], IRAD[45], and AlphaFold-Multimer[46], reveals that while reported success rates for general PPI predictions range from 35% to 90%,[43, 44] the performance drops considerably for Ab-Ag interactions, with success rates between 19% and 63%[43, 46]. This disparity highlights the limitations of existing models and underscores the need for dedicated structural datasets, such as VASCO, to enable the development of machine learning and computational approaches specifically tailored for antibody-antigen interactions.

The key findings from the dataset analysis are as follows. The pairwise contact sequence signature in viral Ag-Ab interactions is distinct from general GPPI, reflecting the specialized nature of immune recognition. Secondary structure analysis indicates not only disordered coils and turns, but also a preference for β-bridges in antigen binding regions, contrasting with the more diverse structural motifs in GPPI. Manifold embedding reveals consistent patterns across viral species, clustering distinctly from GPPI that shows greater divergence, highlighting fundamental differences in interaction landscapes. Interestingly, HIV exhibits a unique interaction pattern, likely due to its heavily glycosylated envelope, which constrains accessible antibody-binding sites.

Moving forward, the VASCO dataset offers numerous opportunities for research and application. As researchers develop more sophisticated machine learning algorithms for antibody-antigen interaction prediction, this dataset will be instrumental in training and testing these models. By integrating data-driven approaches with the structural features highlighted in this study, future predictive tools will be better equipped to model the complexities of viral immunity. Moreover, the dataset’s focus on viral interactions provides an invaluable resource for pandemic readiness. By expanding our understanding of how antibodies interact with viral antigens, we can better anticipate viral mutation impacts, refine therapeutic interventions, and prepare for emerging viral threats.

## AUTHOR CONTRIBUTIONS

S.C. conceptualized the study. K.W., I.S., E.S. and S.C designed the experiments. K.W., I.S., E.S. and S.C performed the data collection, analysis, interpretation, and figure preparation. K.W., I.S., E.S. and S.C wrote and reviewed the manuscript. S.C. is the corresponding author.

## COMPETING INERESTS

The authors declare no competing interests.

## DATA AND MATERIALS AVAILABILITY

Data and materials are available from the corresponding authors upon request.

