## Supplementary material for "Unveiling Interaction Signatures Across Viral Pathogens through VASCO: Viral Antigen-Antibody Structural COmplex dataset": Methods and Supplemental Information

### METHOD DETAILS

***Data Curation:*** We exhaustively searched for protein structures on the Protein Databank[1] with both viral antigen and antibody components, accounted for redundancy and repeats. We noted the chains that make up the structure and disregarded structures with resolutions worse than 5Å. The control dataset of both heterodimer and homodimer general PPIs was curated from the DOCKGROUND dataset[2]. Our dataset consists of 1225 viral Ab-Ag interactions, 2000 heterodimers, 2000 homodimers.

***Interface Extraction:*** The 20Å Ab-Ag interface region is defined as the union of the antigen region within 20Å of the antibody and the antibody region within 20Å the antigen region. These interface regions were extracted using VMD 1.9.4[3].

***Residual Contact analysis:*** We first parsed the interface PDB file using the BioPython.PDB module[4] then defined residual contacts as residues between antibody and antigen chains with heavy atoms within 4.5Å of each other.

***Hydrogen Bonding:*** PyMOL 2.5[5] was used to add hydrogen atoms to the interfaces as hydrogens are typically not considered in experimentally determined protein structures from x-ray crystallography or electron microscopy. Then, VMD 1.9.4 was used to extract the number of hydrogen bonds between the antibody and antigen chains. Hydrogen bonds were assumed when a donor and acceptor on interacting proteins were within 3.5Å and the angle between the donor, hydrogen, and acceptor was less than 60°.

***Secondary Structure Composition:*** We determined the relative frequencies of secondary structures for 10Å interface regions using the STRIDE algorithm[6] implemented in VMD 1.9.4. The secondary structures assessed include coils, turns, beta sheets, isolated bridges, alpha helices, 3-10 helices, and pi helices.

***Binding Region Surface Area:*** Solvent accessible surface areas (SASA) of PDB structures were calculated using the Shrake-Rupley algorithm[7] with a 1.4Å. The contact surface area is calculated as follows: $[A_{ab} + A_{ag} - A_{ab:ag}]/2$ where $A_{ab}$ and $A_{ag}$ denotes the SASA of antibody and antigen chains respectively and $A_{ab:ag}$ is the SASA of the entire complex.

***Sequence-based Contact Maps:*** We first parsed the interface PDB file using the Bio.PDB module and defined residual contacts as residues on antibody and antigen chains with heavy atoms within 4.5Å of each other. Any heavy atom on an antigen chain residue within 4.5Å of any heavy atom on the antibody chain residue qualified as a "contact." For a given PDB file, each antigen chain was compared to the heavy chain and light chain of the antibody. Nonstandard amino acids were skipped. This process was done for every viral Ab-Ag structure that was collected. For the dimeric structures in the control sets, the two chains were simply compared to each other. Contact results were written to text files and visualized on frequency heatmaps generated using the matplotlib.pyplot module[8].

***Structural Energy Minimization:*** Before carrying out the binding energy calculation, each PDB structure had to be energy-minimized. First, the PDB file was centered and the required protein chains were isolated using TCL scripts with the Visual Molecular Dynamics (VMD) program[3]. Missing heavy atoms (where present) in the structure were also added using PDBfixer from OpenMM[9]. After this, a series of GROMACS (version 5.1.4)[10, 11] commands were executed to generate a suitable input file for energy minimization (EM). The pdb2gmx command was used first to generate a .gro (GROMACS file format) coordinate file, using the CHARMM36m[12] forcefield and the TIP3P water model[13]. A new PDB file was also made using pdb2gmx using the same forcefield and water model. This "processed" PDB file was used as the input for gmx make_ndx, in order to create a detailed index file for later use with g_mmpbsa. This index file is important to differentiate the antigen and antibody in the binding energy calculation, since .gro files do not support chain identifiers on their own. In the index file, the antigen chains were collectively labeled as the ligand while the antibody chains were labeled as the receptor.

The structure (.gro file) was then simulated in a cubic box using gmx editconf, with the structure centered inside and placed at least 1.0 nm from the edges of the box. Once the box was defined, it was filled with water using the gmx solvate command. The spc216.gro solvent configuration was used, which is a generic equilibrated 3-point solvent model that is compatible with the TIP3P water model[13] and comes standard with GROMACS installations. Ions were added next using the grompp and genion commands. Grompp was used to generate an atomic level input (.tpr) file from a separate .mdp file that describes standard parameters of the system. Once the .tpr file was generated, it was used to add potassium and chloride ions with genion, specifically by replacing random water molecules in the box with ions. The protein was neutralized with an ion concentration of 0.15M. With the neutralized protein, grompp was used again to assemble the structure, topology, and simulation parameters into a single .tpr file. Finally, energy minimization was carried out by invoking mdrun. This generates four files: a .gro energy minimized structure, a trajectory (.tpr) file, an energy file, and a text log file of the energy minimization process.

***Binding Energy Estimations:*** The binding energy of our complexes was calculated using the Molecular Mechanics Poisson-Boltzmann Surface Area (MM-PBSA) method from the g_mmpbsa package[14, 15]. The package is based on GROMACS and APBS and provides many options for selecting atomic radii and different solvation models. With the energy minimized structure, g_mmpbsa was used to calculate the binding energy between the antigen and antibody. Four input files were passed to g_mmpbsa: the EM .gro structure, the EM .tpr file, a .mdp parameter file, and the index file generated earlier for the structure. The parameter file was provided by the g_mmpbsa GitHub [4]. The solute dielectric constant of 4.0 was used, and the non-linear Poisson-Boltzmann equation was chosen to be solved. The nonpolar solvation energy model was described as a combination of solvent accessible surface area (SASA) and solvent accessible volume (SAV) functions.

The entire EM and MM-PBSA process was performed for all collected viral and control structures. Energy values for the total binding and each energy subcategory were read and stored. Plots were constructed using matplotlib.pyplot[8] for each energy category to identify differences between the Ab-Ag complexes and the control structures.

***Dimensionality Reduction:*** Using dimensionality reduction algorithms to analyze molecular dynamics (MD) trajectories has become commonplace as it allows researchers to focus on the main conformational changes of the trajectory and filter out minor changes by taking the position of each atom in the trajectory as a dimension[16]. Treating our dataset as a trajectory from an MD simulation allows us to use those same algorithms. We start by simplifying each residue in the interfaces down to only central-alpha-carbon atoms (CA) and standardized the atom count of each interface, keeping the 100 closest CA atoms from the antibody chains closest to the antigen chains and vice versa. One of the pdb files (PDB code 8SAV) was randomly designated as a 'reference structure' to remove all translational and rotational deviations between different complexes. The dimensionality reduction algorithms of principal component analysis, isomap, and spectral embedding were performed using the scikit-learn package[17]. Amongst the dimensionality reduction methods, there really is no "best" method and different techniues can work better in different problems, depending on the higher dimensional manifold shape[16].

***Extraction of representative structures from the embedded extremes:*** Spectral embedding showed interesting clustering when plotting dimension 1 against dimension 2. There were three areas on the plot which showed a high density of structures, so they were investigated further. The goal was to use gmx cluster [9, 10] to find a representative viral and control structure for each of the three areas.

Numerical bounds for each high-density area were defined and the specific coordinates for every structure within the bounds was identified for each area. Each coordinate corresponded to a 200-ramino-acid interface that we isolated from the collected PDB files. The interfaces were concatenated to create a single file with all of the structures in the given area. Viral and control structures were separated, resulting in six total concatenations (2 per area).

To create the trajectory file, gmx trjconv [9, 10] was used. The first frame in the concatenated structure file was used as the reference structure and the selected fit was rot+trans (rotation and translation). Then, gmx cluster[10, 11] was performed to identify a single "cluster" of structures. The linkage method was used with a cutoff distance of 1 nm, meaning two structures have to be less than 1 nm apart to be considered neighbors. Least squares fitting was also used before RMSD calculation (-fit option). 1 nm was used as the default cutoff and raised slowly (3, 5, 10, etc.) until a single cluster was found. This resulted in a PDB file containing the mean structure for the area and group (viral or control) of interest.

The cluster file and the trajectory were then loaded in VMD. The RMSD trajectory tool was used to find the representative structure. For each cluster, the trajectory file (from trjconv) and cluster file were loaded into the tool, using the cluster file as the reference. The trajectory was aligned to the reference and the RMSD calculation was performed. The trajectory frame with an RMSD value closest to zero was identified as the representative structure. This frame was cross-referenced back to the structure interfaces we collected to identify the real representative structure for each area of interest on the spectral embedding plot.

### SUPPLEMENTARY FIGURES

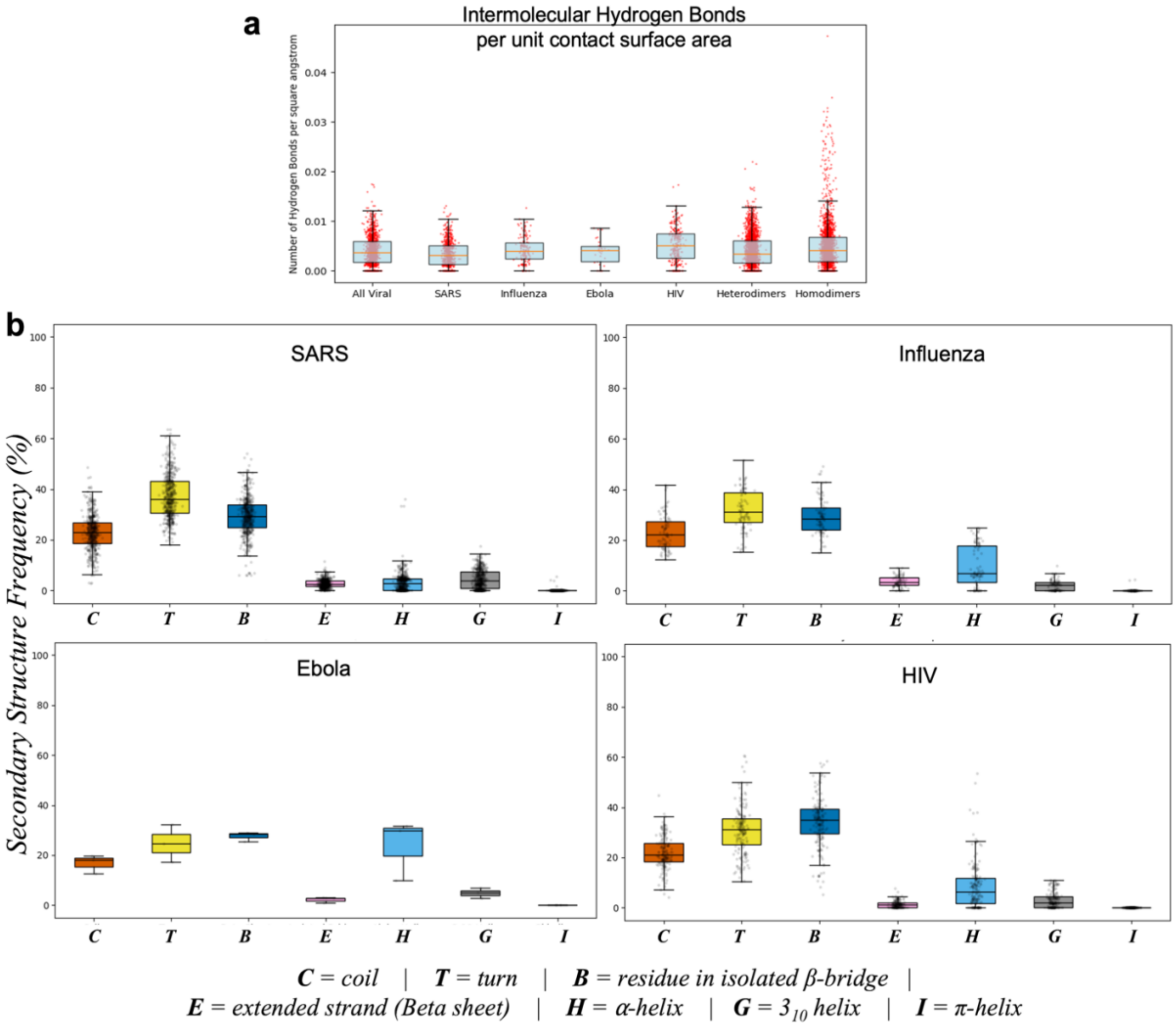


**Figure S1: Structural characterization of VASCO and its comparison to the GPPI heteromeric and homomeric complexes**. (a) Fractional count of intermolecular hydrogen bonds between participating proteins, per unit surface area of contact. Median and quartile ranges are shown in the box plots.(b) Median and quartile ranges of secondary structure distributions over interface residues for SARS, Influenza, Ebola and HIV interfaces within the VASCO set.

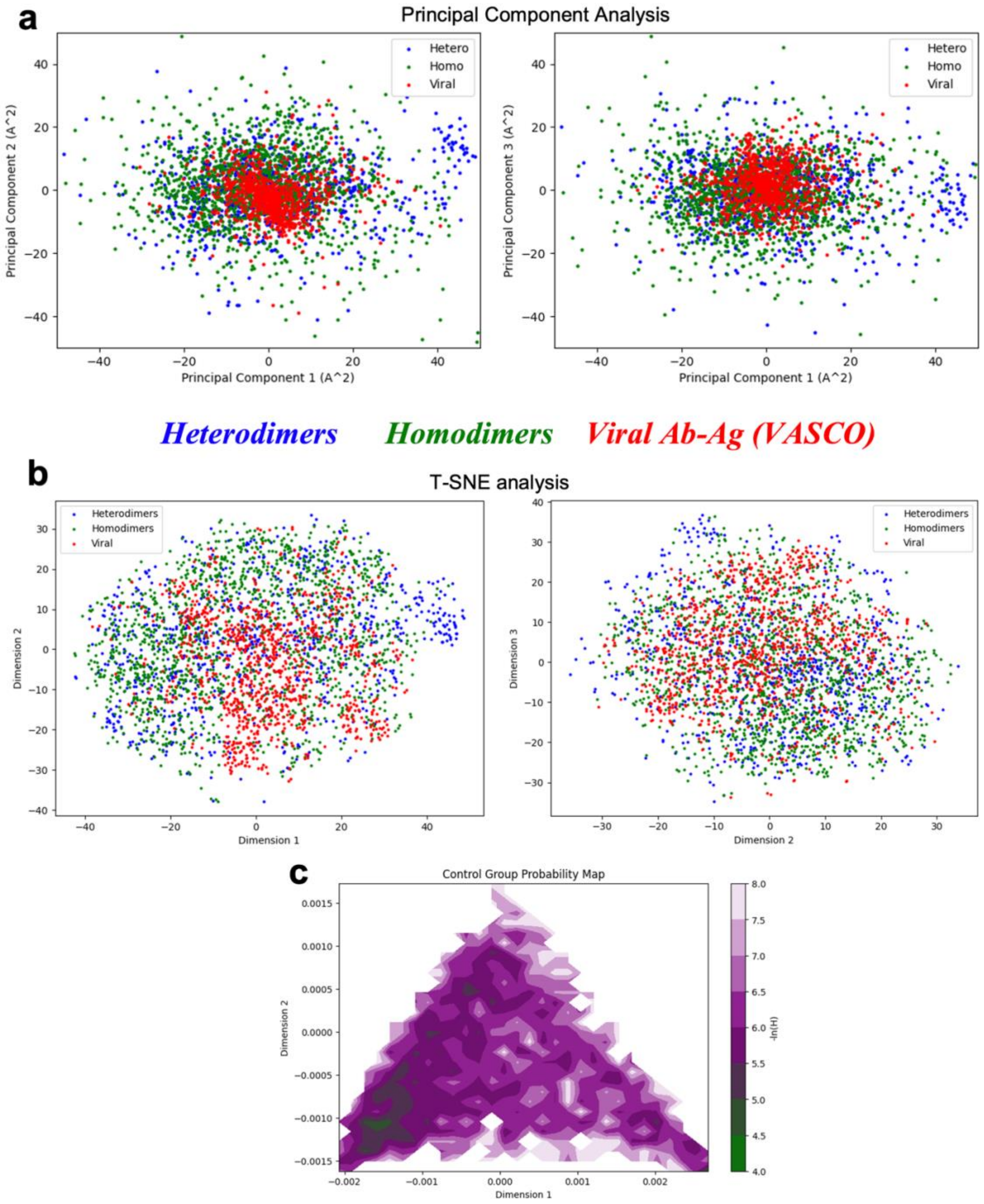


**Figure S2: Manifold reduction analysis of viral Ag-Ab and general PPIs**. (a) Principal Component Analysis with PC 1 and PC 2 dimension projections and (b) PC2 and PC3 dimension projections of the combined set of VASCO, and general protein-protein interface structures. The top three highest PCs could not capture differences between VASCO and GPPIs. (c) Negative log probability map of an interface, measuring its likelihood of being a general PPI (and not a viral Ag-Ab complex).
